# Methods for the analysis of skin microbiomes: a comparison of sampling processes and 16S rRNA hypervariable regions

**DOI:** 10.1101/2023.01.28.525888

**Authors:** R. Y. Alyami, D. W. Cleary, J. Forster, M. Feelisch, M. R. Ardern-Jones

## Abstract

**Background:** The skin microbiome is increasingly recognised as a critical component of the innate skin immune response and is central to the pathogenesis of many inflammatory skin disorders including atopic dermatitis. Previous studies have not looked in detail at the impact of changing both sampling methodology and hypervariable region sequencing for skin microbiome analysis.

**Objectives:** We set out to undertake a detailed analysis comparing microbe population diversity and resolution at the single species level by swab, tape, scrape and scrape then swab sampling assayed by hypervariable region sequencing.

**Methods:** In triplicate samples from the antecubital fossa were taken from healthy volunteers for 16s RNA analysis by primer amplification for hypervariable regions 1-3, and 3-4.

**Results:** Alpha (phylogenetic) diversity was the greatest with tape sampling and V1-3 analysis, whereas for V4, tape and swab were equivalent. Scrape sampling showed lower alpha diversity at both V1-3 and V4. All measures of beta diversity showed the scrape methodology yielded a lesser diversity than the others. Phyla composition was similar across all sampling methodologies. Minor differences in composition were noted between V1-3 and V4 sequencing, but V1-3 was optimal for identification of firmicutes (including staphylococci).

**Conclusions:** In the methodological planning of skin microbiome analysis skin scientists need to consider the microbes of interest to choose the optimal hypervariable region to sequence, and harmonisation of methodological approaches would be beneficial to the field. For detection of staphylococci on flexural skin, we would recommend tape sampling with analysis of V1-3.

**What’s already known about this topic?:**

- Previous microbiome studies have utilised many different methodologies, but a detailed comparison of their utility in skin research has not been undertaken
- Methodological choices in skin microbiome analysis affect both sensitivity to detect diversity and sensitivity to identify at a species level
- Optimisation to detect diversity and species identification can require different approaches

**What does this study add?:**

- Our study compared a matrix the four most widely utilised skin sampling methodologies and the two principle means for hypervariable gene analysis.
- We show that alpha diversity is optimal with tape sampling
- Additionally, in contrast to gut microbiome analyses, in the skin hypervariable region V1-3 analysis is superior to V4 when the species of interest are staphylococci.

## Introduction

The skin is among the largest organs of the human body and is populated by a diverse array of microorganisms (1). There is increasing evidence that the skin microbiome plays a crucial role in the defence against pathogens, the education of the immune system, and skin homoeostasis (1, 2). Unsurprisingly, shifts in skin microbiota composition are associated with skin disorders like acne, atopic dermatitis, eczema, and psoriasis (3). For example, the presence and/or abundance of pathobionts such as *Staphylococcus aureus* have been shown to correlate with disease severity of atopic dermatitis (3).

To better understand the microbiome-host crosstalk in the context of specific cutaneous diseases, and as an avenue to assess potential intervention strategies, it is important to develop reliable tools for characterisation of the composition of the skin microbiome beyond the limitations of culture. Whilst several molecular profiling approaches based on high-throughput sequencing of microbial 16S ribosomal RNA have proved successful in addressing this (4, 5), specific technical challenges remain such as low microbial biomass and the impact of extensive background human DNA material that have implications for choosing the most adequate method to reconstruct the microbial community composition.

The first decision investigators have to make is to select the most appropriate sampling methodology. Although, arguably, skin biopsy would be ideal and allow further spatial selection of compartments within skin (e.g. using laser-capture microscopy), it is an invasive procedure, complicating sampling from multiple body regions of the same individual (6, 7). Whilst previous reports of sampling using pre-moistened swabbing (4, 8, 9) suggest that the relative proportions of bacterial species this technique recovers are comparable to a skin biopsy (6), alternative methods for skin microbiome sampling may be preferable to wet swabbing. Moreover, it is challenging to minimise methodological variation with wet swabbing for quantitative analyses. For example, it is difficult to control for swab pressure, frequency and direction of swabbing (10). In contrast, tape adhesion sampling, is a straightforward technique for investigators, which yields high sample biomass and uniformly samples a specific surface area of the skin (11, 12). Skin scrape methods further increase the captured biomass and are therefore likely to be useful for lower abundance microorganisms (6, 13). Cup scrubbing and cylinder suspension are alternative sampling techniques that have been reported to yield higher levels of viable bacteria from a well-defined surface area compared to traditional swabbing methods (14, 15).

The second decision to make before engaging in a skin microbiome study relates to the molecular methodology employed for sequencing. One of the main challenges for taxonomic analysis of 16S rRNA sequencing data is the relatively short read lengths (up to 300bp for single-end and 600bp for paired-end) obtained from the most commonly employed next-generation sequencing approaches. Thus, sequence resolution down to species level is enhanced by choosing the most appropriate hypervariable region of the 16S rRNA gene relevant to the expected microbiota in the sampled region and is therefore to some degree dependent upon the tissue of interest. In the skin, differentiation within a genus is critically important because of the recognised differences between microbial species. For example, within staphylococci, distinguishing between homeostasis regulating *S. epidermidis* and inflammation inducing *S. aureus (SA)*is important for many studies (16). Of the nine 16S rRNA gene hypervariable regions (19), V1 to V9, region four (V4) has been used most widely in studying human gastrointestinal tract microbiome. However, recent work has shown that 16S rRNA V1-3 region demonstrate a good taxonomic classification result including an increased resolution for *Staphylococcus’* species (17). Furthermore, V1-3 has the advantage of showing similar skin microbial communities and genetic functional profiles to whole metagenome shotgun (WMS) sequencing result as compared to V4 region (5, 18) and is more sensitive at detecting other skin microbiota (e.g. Propionibacterium)(19).

Previous analysis using V4 sequencing of skin has shown good concordance between swabbing, tape stripping and cup scrubs (12), but because different biomass yields are returned by the various sampling methodologies, it is likely that the choice of sampling will interact with the choice of 16S rRNA sequencing target. As a result, detailed comparisons between sampling methodology and 16S rRNA target are important.

The aim of the present study was to compare microbiome composition using two 16S rRNA hypervariable molecular targets across sampling techniques, to select the optimal methodology for identification of skin microbial diversity.

## Methods

### Ethics

Ethical approval was obtained from the Northern Ireland regional ethics committee (NRES 15/NI/0180). Local ethical approval was granted by the University of Southampton (ERGO ID 17473) and the University Hospital Southampton NHS Foundation Trust (R&D CRI 0320).

### Skin Microbiota Sampling

Four healthy volunteers (male, aged 26-40) were recruited for this study. Four methodologies, each performed in triplicate, were compared for sampling from the same antecubital fossa. Relevant control samples were analysed for each sampling method. Prior to sampling, the investigator’s gloves were treated with 70% v/v EtOH and air dried prior to contact with the subject.

Method 1 - Swab: Tubed sterile dry tipped swabs™ (MWE medical wire^®^, UK), coated in 0.15 M NaCl + 0.1% Tween20 (SigmaAldrich, UK) were turned back and forth over a defined 4 cm² area of skin for 30 secs.

Method 2 - Scrape: A 4 cm^2^ area of skin adjacent to the swabbed region was outlined and then scraped gently with a surgical scalpel blade (No.15, Williams Medical, UK) with the blade curve horizontal to the skin surface to obtain microbial DNA(9). Materials were collected from the blade using sterile dry swab tips (MWE medical wire^®^, UK), coated in 0.15 M NaCl + 0.1% Tween20.

Method 3 - Swab after scrape: Following the scrape (Method 2), the same area was swabbed exactly as described in Method 1.

Method 4 – Tape adhesion: Sampling tape (D-Squame, CuDerm Corporation) was sterilised by submersion in 70% v/v EtOH. The adhesive portion of the 4cm^2^ tape disc was placed on the surface of the skin and even pressure applied down with the gloved thumb. The disc was then peeled off the skin and the same single tape disc was reapplied to the same area of skin 50 times over a period of 2 min.

### DNA Extraction

Sterile forceps were used to place swab tip or sealed tape into a PowerBead^®^ DNA extraction tube (Qiagen, UK). Total DNA extraction was accomplished using PowerSoil^®^ DNA Isolation Kit (Qiagen, UK) according to the manufacturer’s instructions.

### 16S rRNA PCR

The 16S rRNA PCR was done using primers 27F (5ʹ-AGAGTTTGATCMTGGCTCAG-3ʹ) and 534R (5ʹ-GTGCCAGCAGCCGCGGTAA-3ʹ) for variable regions 1 to 3 (V1-3) and 515F (5ʹ-GTGCCAGCMGCCGCGGTAA-3ʹ) and 806R (5’-GGACTACCGGGGTATCT-3’) for V4 (20). PCR was performed using a GeneAmp^®^ PCR System 9700 (Applied Biosystems, UK) with the following cycling conditions: 3 min of denaturation at 98°C followed by 35 cycles of 98°C for 30 secs, 55°C of annealing for 30 secs, and extension at 72°C for 30 secs, followed by a final extension at 72°C for 5 min. Each sample was amplified in triplicate 25 μl reactions consisting of primers at final concentrations of 0.2 μM and 1X PCR reagent (Q5^®^ Hot Start High-Fidelity 2X Master Mix, New England BioLabs, UK). Triplicate PCR reactions for each sample were then pooled and purified using Agencourt^®^ AMPure^®^ XP beads (#A63881, Beckman Coulter, UK).

### Library Preparation

Adapter ligation and barcoding were performed using the Nextera XT Index Kit (#FC-131-1002, Illumina**^®^**, UK). PCR conditions involved 3 min of denaturation at 98°C, then 12 cycles at 98°C for 30 secs, 55°C of annealing for 30 secs, 30 secs of extension at 72°C, followed by a final extension at 72°C for 5 minutes. The number of PCR cycles was kept low to reduce amplification bias, and reduce the risk of amplifying contaminants present in kit reagents. This PCR amplification was followed by another purification using the Agencourt^®^ AMPure^®^ XP beads (#A63881, Beckman Coulter, UK). A ratio of 6.5 of Solid Phase Reversible Immobilisation beads (SPRI) to DNA was used for size selection. The library was quantified using a Qubit™ double stranded DNA (High Sensitivity) Assay Kit (#Q32866, Life Technologies Ltd, UK) with the Qubit^®^ 2.0 Fluorometer (Life Technologies Ltd, UK).

### Sequencing

The DNA library sample pool (2nM) was denatured using 0.2N NaOH, diluted to 10pM, and combined with 25% denatured 4nM PhiX. Samples were sequenced locally (within the Wessex Investigational Sciences Hub (WISH) facility, University Hospital Southampton NHS Trust) using a MiSeq (Illumina^®^, UK) to generate 2 × 300 bp paired-end reads using V3 chemistry, following standard sequencing protocols.

### Analysis

Initial processing and analysis was performed using the Quantitative Insights in Microbial Ecology (QIIME2) package (21). Individual forward and reverse read files were imported into QIIME2 as a single merged file using the provided import tool. Reads were demultiplexed and the trimmed using a quality score threshold of <Q35. Only forward reads were processed due due to low quality of reverse reads. After trimming, the reads were denoised using DADA2 (22), exact amplicon sequence variants (ASVs) inferred, aligned and finally given a taxonomic label using the Greengenes database (release 2013-08; gg_13_8_otus). The output QIIME artefacts, namely the feature table, taxonomy, phylogenetic tree and metadata, were then used to create a phyloseq object using qza_to_phyloseq from the package qiime2R (23)(v 0.99.12). Subsequent analysis was done using phyloseq (24) (v 1.28.0). Contaminant ASVs were identified in the controls and excluded using the R package ‘decontam’ (25). All analysis was done in R (v 3.6.1) using RStudio (v 1.2.5019).

### Statistical Analysis

Alpha diversity were tested using (one-way ANOVA, p> 0.05) and beta dissimilarity score tested using PERMANOVA test, p> 0.05) in R (v 3.6.1) using RStudio (v 1.2.5019). Figures were produced using the package gplot2 (v.3.3.3).

Further analysis was conducted using GraphPAD Prism^®^ software (v9, GraphPad Software, Inc., La Jolla, CA, USA). Statistical difference was evaluated using paired two-tailed t-test and one-way analysis of variance (one-way ANOVA) test to determine any statistically significant differences between groups and results were considered significant when p<0.05.

## Results

After filtering, the total number of sequences identified across all sampling methodologies was considerably higher for V1-3 (1,848,741) as compared to V4 (627,088). The mean per sample sequence counts excluding negative controls, for V1-3 was 38,515 (range 13,215 to 121,731) vs 13,064 (range 3,990 to 31,786) for V4. In accordance with the higher number of sequence counts, higher numbers of Amplicon Sequence Variants (ASVs) were assigned for V1-3 (2724 taxa) than V4 (1385 taxa) across all sampling methodologies. A two-tailed paired t-test performed on V1-3 and V4 sequence count showed a statistically significant difference (p <0.0001). Despite the low biomass of skin microbiomes and the inherent risks of contamination (26) we observed a negligible amount of contamination, as confirmed by the sequence counts from the negative control samples for V1-3 and V4 (3, 1 respectively).

Alpha diversity (the within-sample ASV richness and evenness) was measured using three separate estimators. The total number of ASVs per sample was significantly higher in samples collected by swabbing, swab after scrape, or tape collection than scrape collection alone (Fig 1a). Similarly, skin microbiome evenness (the relative proportions of identified species, measured using the Shannon index) showed that Scrape sampling identified a less even species distribution as compared to swabbing (p=0.024 V1-3, p = 0.0018 V4, Fig 1b). Community diversity (a quantitative measure accounting for number and abundance of the identified species, measured using Simpsons 1-D index) observed following scrape sampling was significantly lower than both swabbing and swabbing after scraping for V4 (p = 0.0036, p= 0.045 respectively, Fig 1c), but this was not seen with V1-3. Comparison of rarefaction for both regions (Faith-pd) also showed scrape sampling reported the least diverse community, whereas tape sampling was consistently the most diverse (p<0.0001, Supplementary Figure 1). Overall, the V1-3 region showed an approximately 2-fold greater diversity of observed ASVs as compared to V4 with all sampling methodologies (Fig 1a-c).

**Figure 1.**
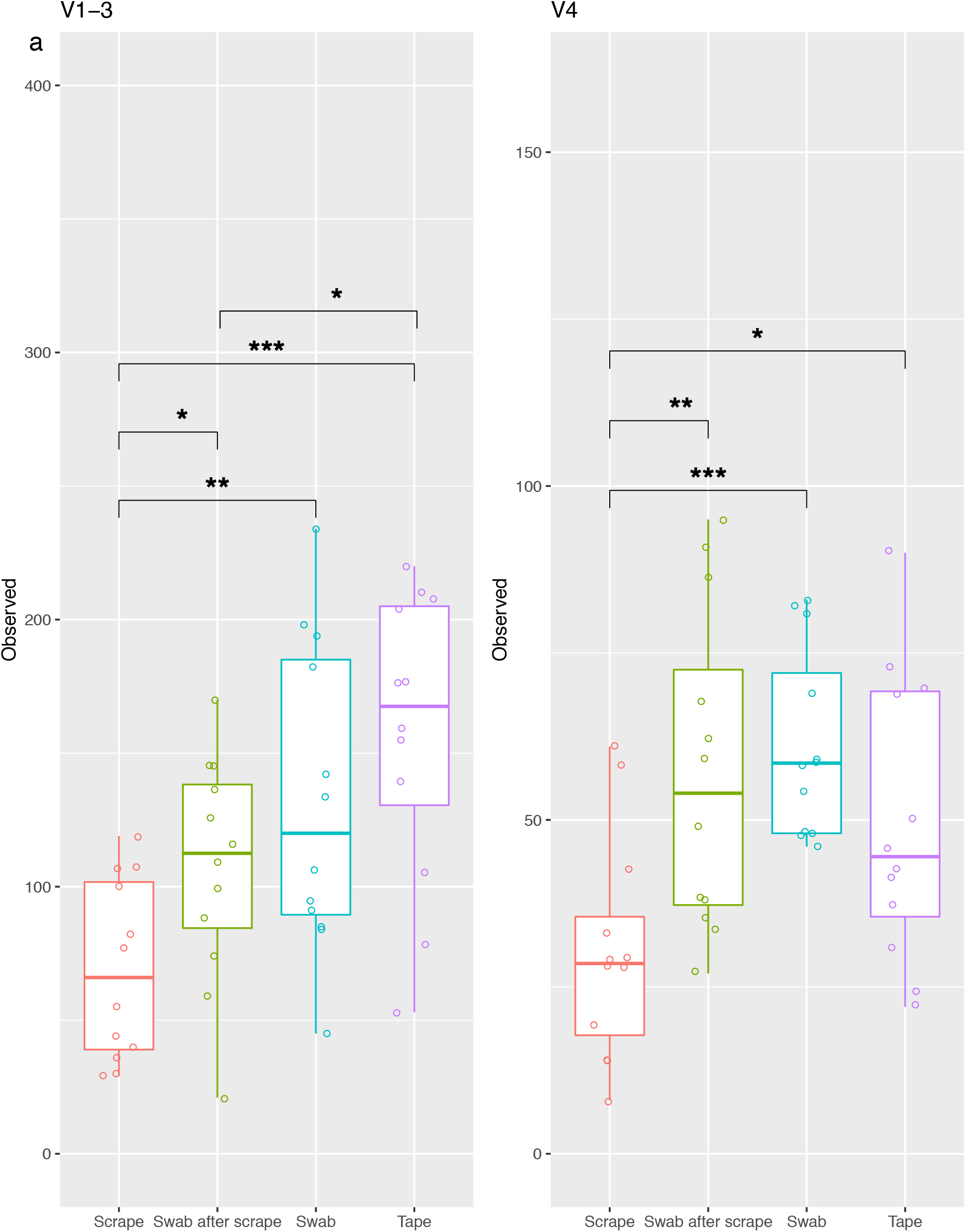
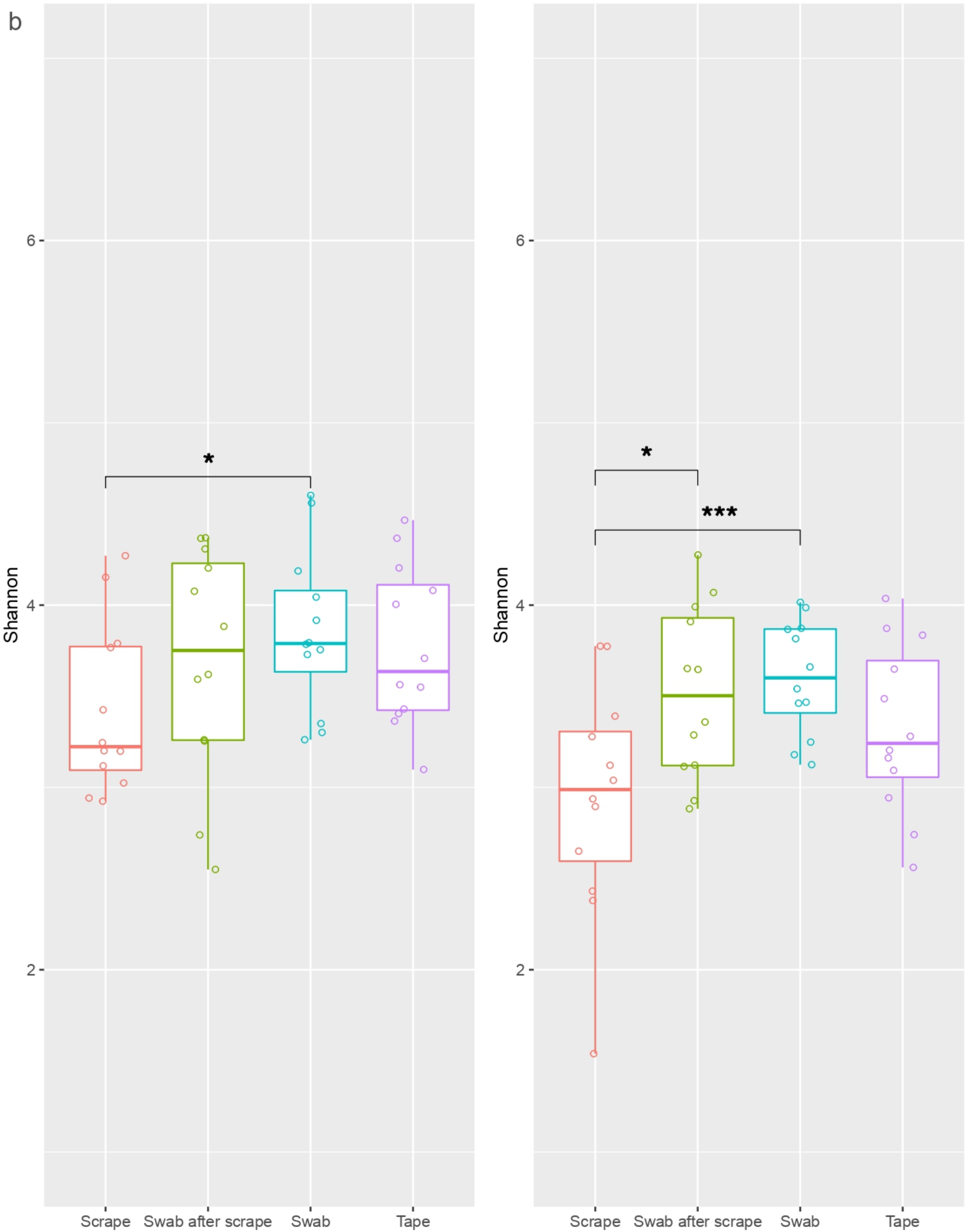
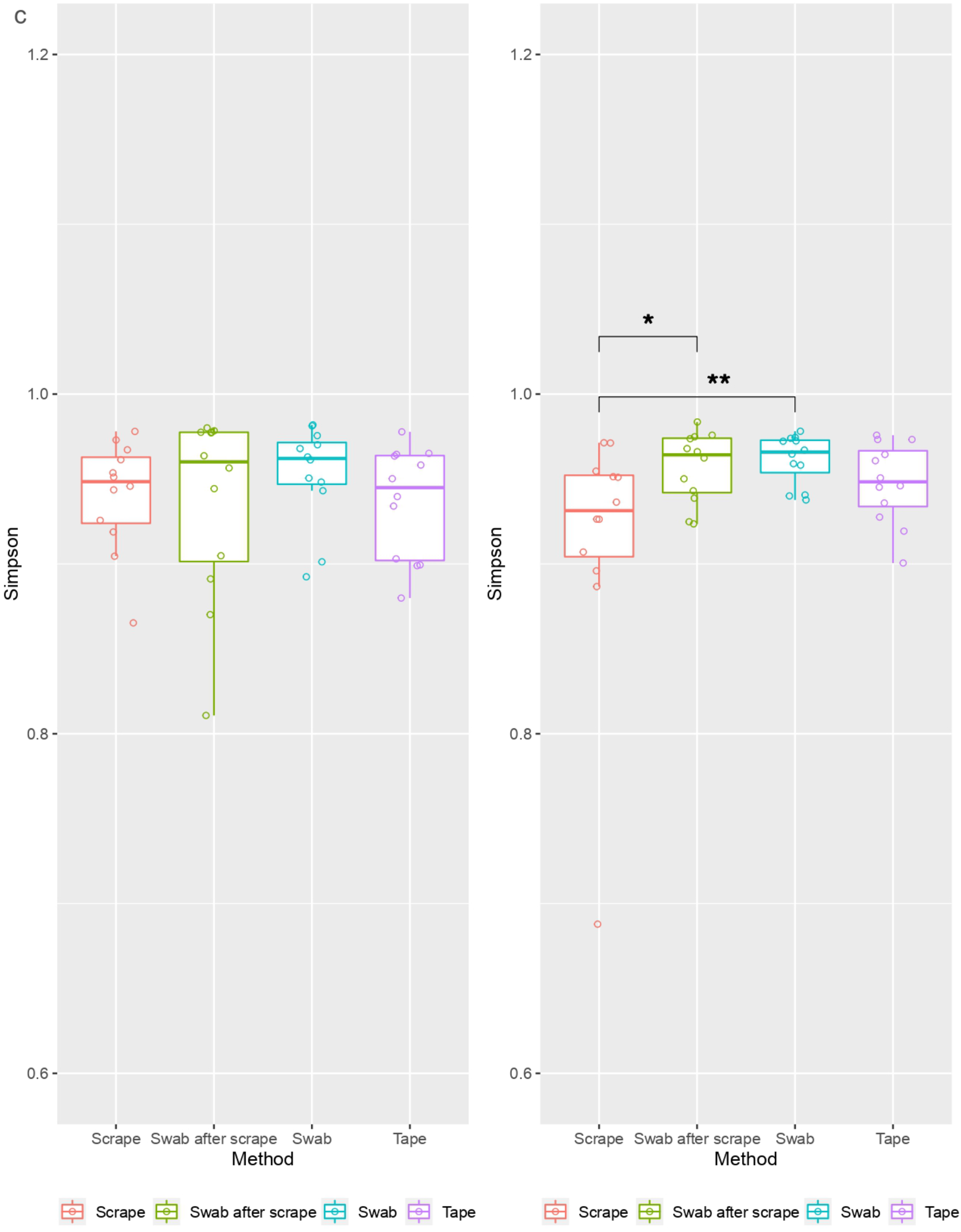
Box and Whisker plot showing alpha-diversity of the microbiomes acquired using four different skin sampling methods from four healthy volunteers. Diversity was estimated using Observed ASVs (a), Shannon (b) and Simpson 1-D (c). Kruskal-Wallis ANOVA for V1-3 (left) and V4 (right) are shown.

To compare compositional differences between samples acquired using the different sampling strategies, principal coordinate analysis (PCoA) of the Bray-Curtis distance dissimilarity score and tested with PERMANOVA (P >0.05) analysis using pairwise adonis tool with 999 permutations (27) (Supplementary Figure 2a, b). There was no statistical difference when swabbing (with or without prior skin scrape) was compared to tape or scrape sampling. However, community composition was significantly different when tape sampling was compared to scrape for both V1-3 and V4 (respectively *p* = 0.009 and *p* = 0.048, Supplementary Figure 2a, b).

Regardless of which 16S rRNA variable region or sampling technique the presence of bacterial phyla comprising *Actinobacteria, Firmicutes, Bacteroidetes, Proteobacteria* and *Fusobacteria*, was generally as expected (Figure 2a, b).

**Figure 2.**
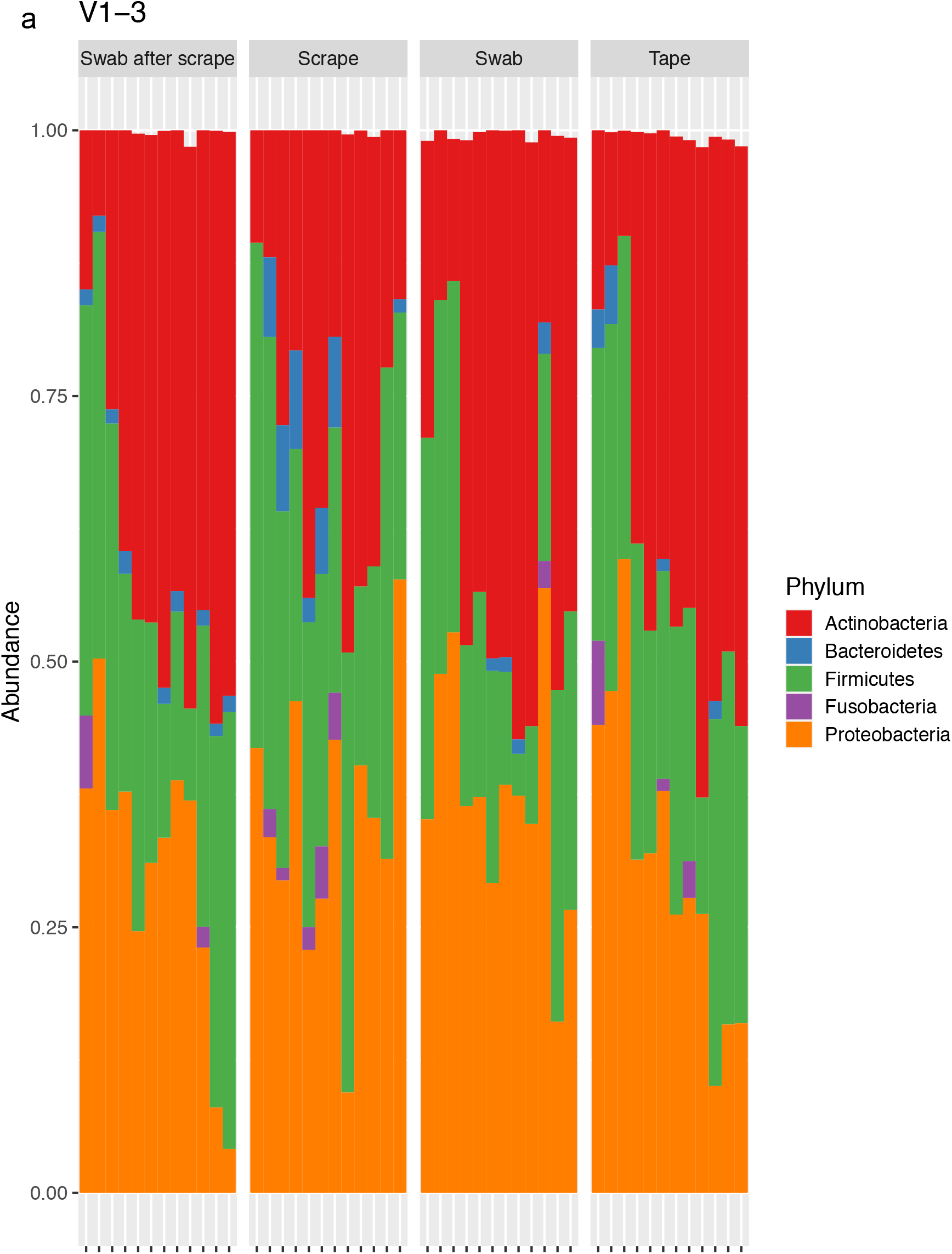
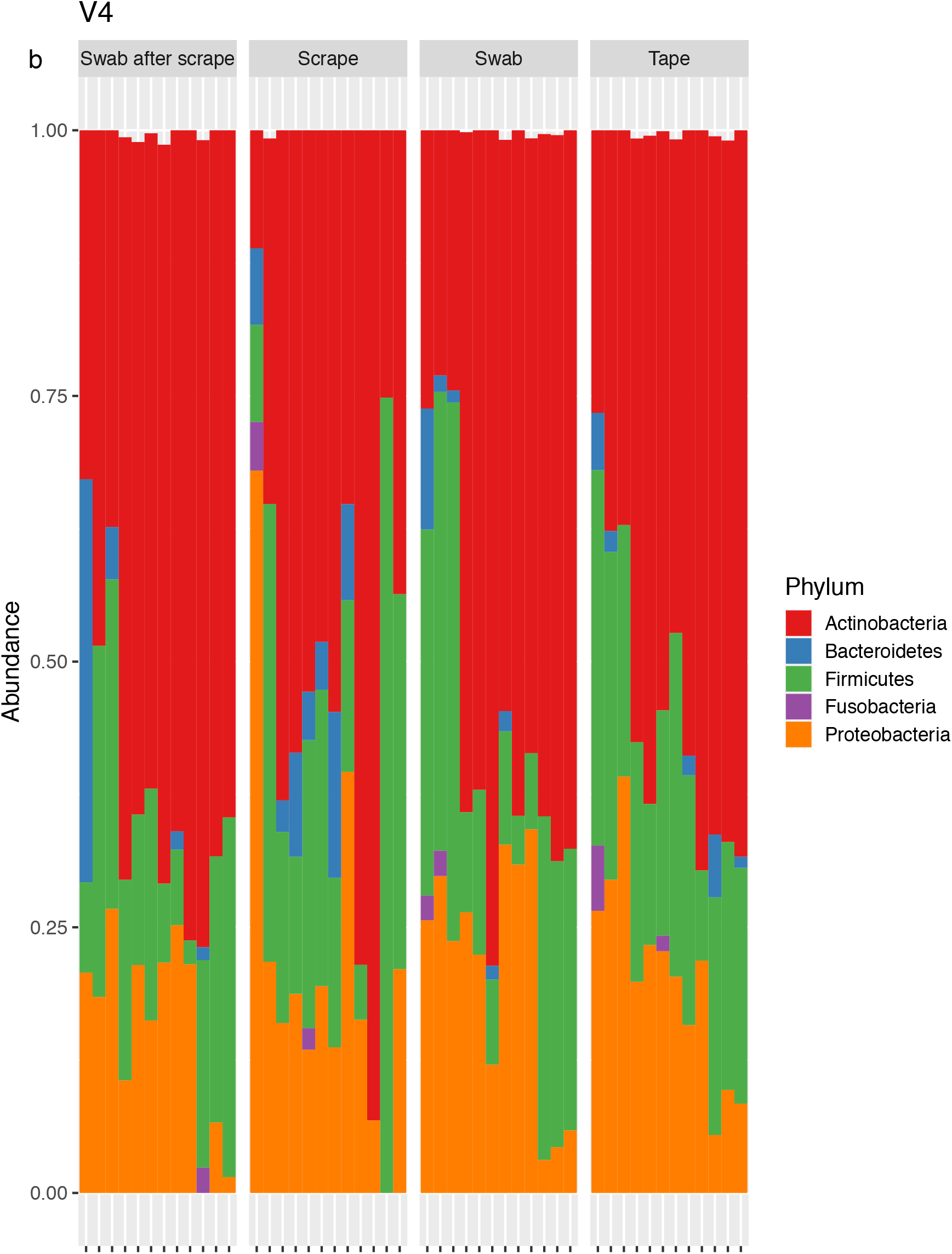
Proportional Bar plots showing the relative abundance of the dominant bacterial phyla using V1-3 across sampling methods. *Actinobacteria* (red), *Bacteriodetes* (blue), *Firmicutes* (green), *Fusobacteria* (purple) and *Proteobacteria* (orange) are shown. (a) V1-3. (b) V4.

Identification of *Bacteroidetes* and *Proteobacteria* was significantly enhanced by scrape sampling as compared to other methodologies on V1-3 analysis (Figure 3). However, this apparent methodological difference was not detected by analysis of V4 (Figure 4).

**Figure 3.**
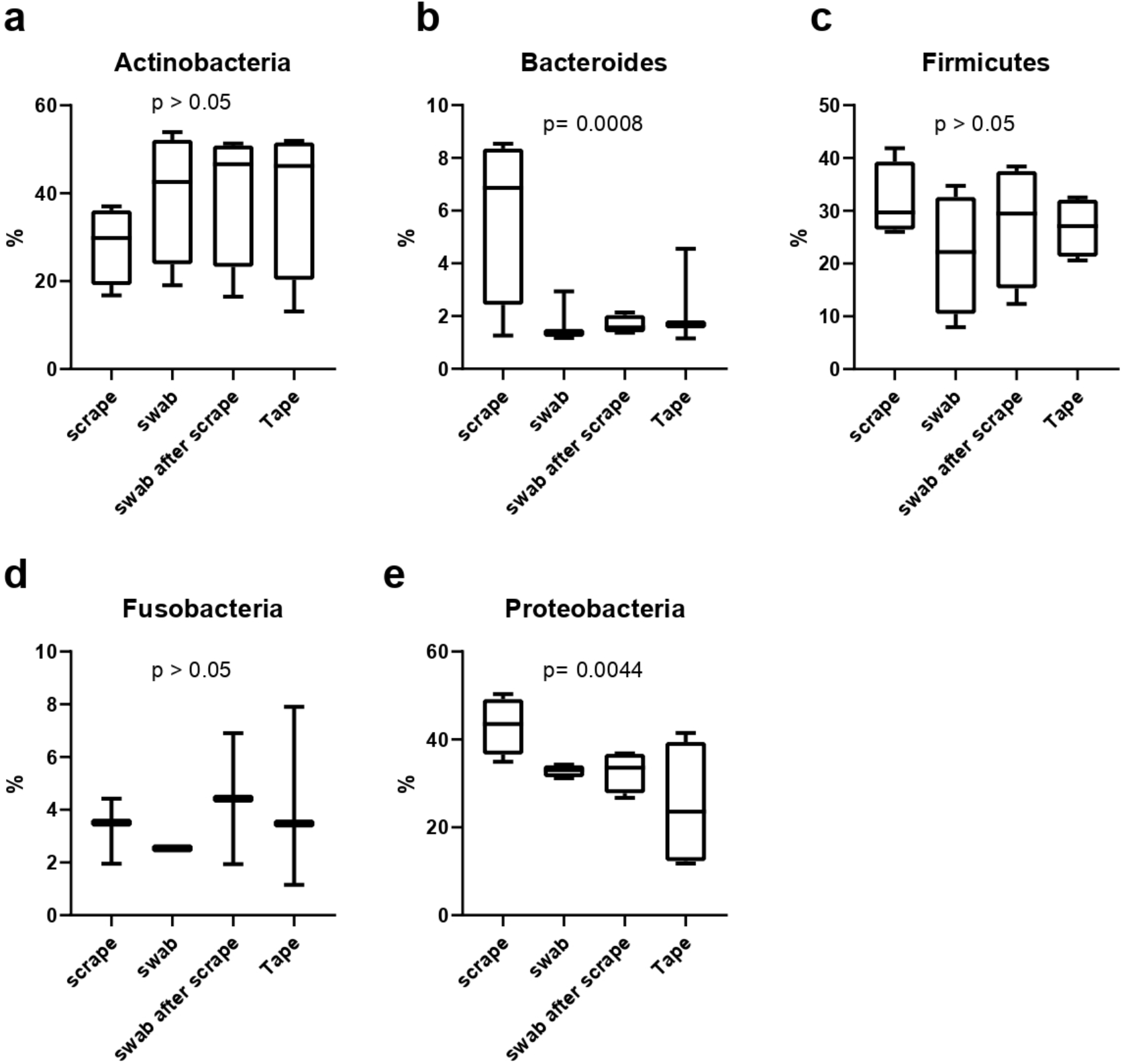
Box plots showing V1-3 relative abundance of identified phyla (a, Actinobacteria; b, Bacteroides; c, Firmicutes; d, Fusobacteria; e, Proteobacteria) by the four used sampling methods: Scrape, Swab, Swab after scrape and Tape.

**Figure 4.**
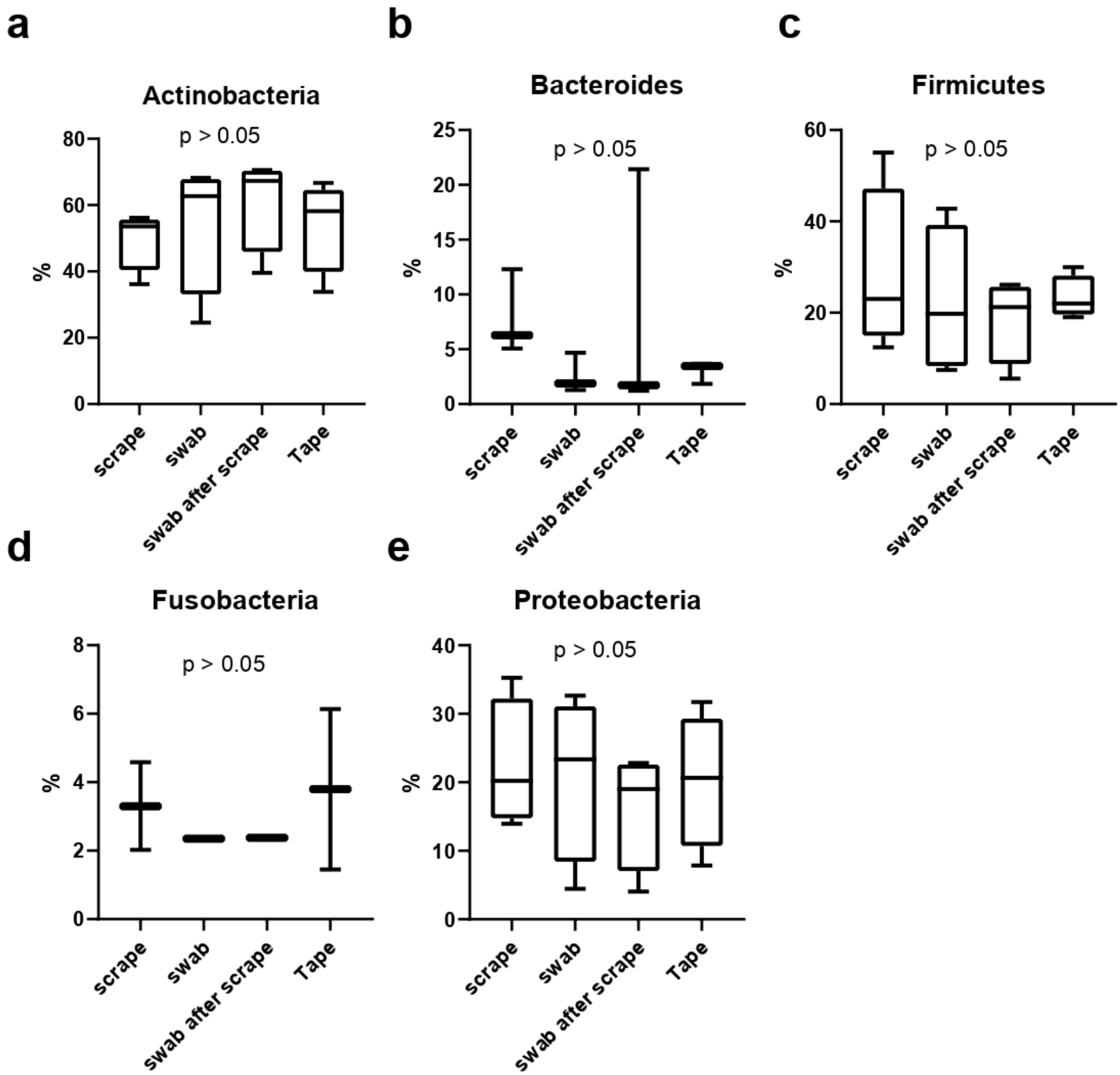
Box plots showing V4 relative abundance of identified phyla (a, Actinobacteria; b, Bacteroides; c, Firmicutes; d, Fusobacteria; e, Proteobacteria) by the four used sampling methods: Scrape, Swab, Swab after scrape and Tape.

Comparing V1-3 and V4 sampling analysis per phyla, V4 was more sensitive in detection of *Actinobacteria* (P<0.05 Scrape, Swab after scrape, and Tape) but resulted in reduced identification of *Firmicutes* and *Proteobacteria* (P<0.05 Swab, Swab after scrape, and Tape) (Figure. 5).

**Figure 5.**
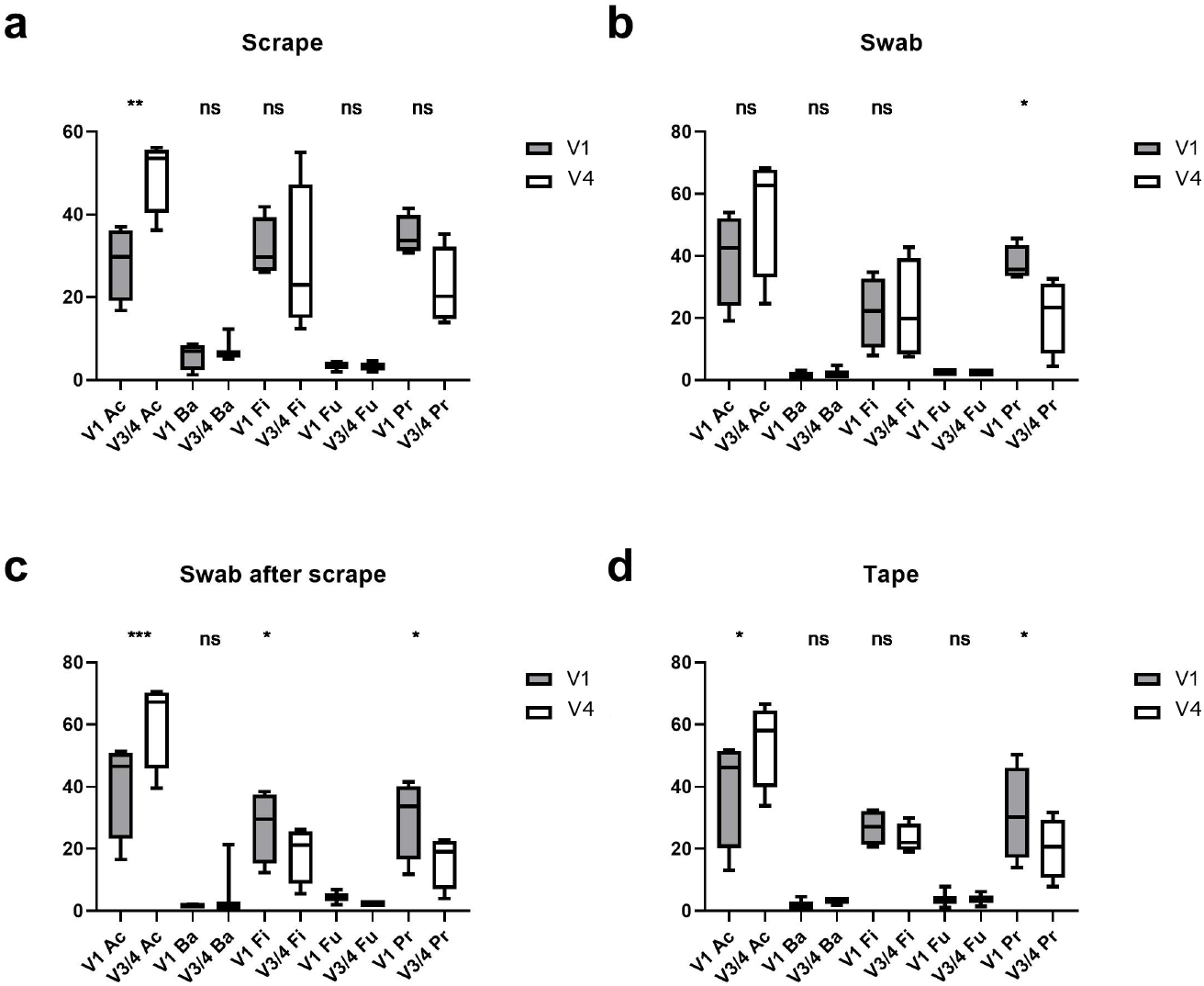
Bar plots showing the relative abundance of identified phyla by V1-3 (grey) or V4(white) analysis. Ac, Actinobacteria; Ba, Bacteroides; Fi, Firmicutes; Fu, Fusobacteria; Pr, Proteobacteria.

Given their importance in skin disease, we next compared V1-3 and V4 for species level identification of *Staphylococcus* sp. (Table 1). Analysis revealed a high proportion of ASVs aligned as belonging to staphylococci. Because *Staphylococcus epidermidis* is ubiquitous in human skin, detection of *S. epidermidis* is expected in all samples. As expected, with V1-3, the dominant staphylococcal species was aligned to *S. epidermidis* with a mean abundance of 18.7-21.5%, and no significant difference between the sampling methodologies was detected (p>0.05, Kruskal-Wallis ANOVA). However, in contrast, no *S. epidermidis* were aligned using V4 analysis. Instead, other staphylococci were classified: *S. equorum* and *S. haemolyticus*. With V1-3 sequencing, S. aureus was identified in two of the four volunteers and where identified the relative abundance of S. aureus was highest in tape samples (3.7%). Whereas targeting V4 detected S. aureus in only one of the four volunteers.

**Table 1:** Relative abundance of Staphylococcal ASVs identified to species level in skin microbiomes using four different skin sampling methods, with V1-3 or V4 16S rRNA analysis. Values represent mean relative abundance with ranges in parentheses.

| **V1-3** | ***S. aureus*** | ***S. epidermidis*** | ***S. haemolyticus*** |
| --- | --- | --- | --- |
| **Tape** | 3.7% | 20.8% (8.8%- 39.7%) | NA |
| **Scrape** | 2.0% | 19.7% (3.8%- 41.7%) | NA |
| **Swab** | NA | 18.7% (4.4%- 36.1%) | NA |
| **Scrape after swab** | NA | 21.5% (1.5%- 39.9%) | NA |
| **V4** | ***S. aureus*** | ***S. equorum*** | ***S. haemolyticus*** |
| **Tape** | 5.5% | NA | 24.4% (7.9% -52.9%) |
| **Scrape** | NA | 8.0% | 10.6% |
| **Swab** | NA | 5.5% | 21.2% (13.0%-37.3%) |
| **Scrape after swab** | NA | 4.2% | 18.9% (5.1%-38.2%) |

## Discussion

Analysis of community compositions in microbiome research is reliant on the use of sampling and analysis methods that give the most accurate representation of the resident microbiota. It is recognised there is no ‘one size fits all’ approach in this context and in this respect the optimal methods the characterisation of human skin microbiota have not been fully determined. Nevertheless, most recent studies used the pre-moistened swabbing technique for sample collection (4, 9, 28). There are various reasons to question whether this is the optimal approach considering, for example, that tape stripping typically yields greater sample biomass and viability (11, 12, 29). Other methods that have been recommended include skin scraping and dry swabbing; while both provide lower biomass, each of them may yield greater specificity (30). Furthermore, whilst recent publications have tended to focus on amplification of the V4 region, direct comparisons of this approach with V1-3 in skin are limited (5, 31, 32).

To determine the optimal sampling method for 16S rRNA hypervariable region analysis we matrixed variable region selection (V1-3 or V4) with replicate sampling methodology. Using Bray-Curtis distance measures to assess beta-diversity, only the scrape samples were significantly different from the other sampling techniques. The data also showed that the scrape method revealed a significantly less diverse and more uneven microbiota composition, suggesting that this approach is inferior for characterising the skin microbiome as compared to the other three methods. Of the other three, swabbing after scrape yielded the most heterogeneous results, but overall, good uniformity between the approaches was detected, with similar measures of skin microbiome evenness and diversity.

Composition of the microbiome in our analyses was broadly as expected for both V1-3 and V4 based on previous work, showing that *Actinobacteria* dominated (51.8%), followed by *Firmicutes* (24.4%), *Proteobacteria* (16.5%) and *Bacteroidetes* (6.3%) (33). However, *Actinobacteria* were identified to a higher abundance with V4 analysis. This might suggest that there is preferential amplification of a member(s) of this Phylum using the V4 primers, but there is little data to support this in the context of skin microbiota (34-36) and we found no clear differences at lower taxonomic levels to support this hypothesis. V4 also showed reduced relative abundance of *Proteobacteria* and importantly *Firmicutes* which includes staphylococcal species.

A greater number of ASVs classified as *Staphylococcus* sp. were identified in V1-3 analysis. Indeed, it is important for skin researchers to note that, in our study of healthy volunteers where S. epidermidis colonisation can be assumed, S. epidermidis was also only identified with V1-3 amplification. It is hard to determine whether this is a feature of the known improvements in taxonomic classification performance of this region for staphylococci (37) or an issue with potential biases in amplification as noted above for V4. Nonetheless, no significant differences between methods were detected for V1-3. The lack of *S. epidermidis* when V4 was used suggests a significant limitation of this approach (38). We did not pre-screen for *S. aureus* carrier status, but the staphylococcal carriage in our population has been previously found to be 28% (95 % CI 26.1–30.2 %) in healthy individuals (39). Here we identified *S. aureus* in 50% of individuals studied with V1-3 analysis, whereas V4 identified *S. aureus* in only 25%. Taken together, this strongly suggests V4 is not suitable if the goal is to detect and distinguish species of staphylococci, a finding that mirrors previous research (5).

Some limitations exist with the study presented here. These include the use of only a small number of healthy volunteers (n = 4), only one body site for sampling and the lack of skin biopsy which was deemed too invasive to justify its inclusion in such a study. Inevitably, this makes extrapolation to different skin types difficult. Additionally, a lack of data on bacterial biomass recovered, which could have been achieved through use of a 16S rRNA qPCR for example, would have added quantitative data and made inter-sampling method comparisons more informative. Pre-screening recruits for *S. aureus* and *S. epidermidis* colonisation would also have been useful to enhance the comparative analysis between sampling approaches and their sensitivity for the detection of these species.

In conclusion, our data provide useful information for investigators in studying the skin microbiome. We show important differences between four commonly used sampling methods, evaluated using 16S rRNA sequencing of the V1-3 and V4 hypervariable regions. To maximise the potential for skin scientists to be able to compare different studies on the microbiome, it is important that technical aspects of the methodology are harmonised as far as possible. We suggest that the optimal approach to characterising diversity, with potential advantages for the recovery of *S. aureus*, incorporates tape or swab sampling followed by sequencing analysis of the V1-3 16S rRNA hypervariable region.

## Acknowledgments

We thank all volunteers for their time and involvement in this study. We also thank the research nursing staff at the Wellcome Clinical Research Facility for their support.

## Figure Legends

Supplementary Figure 1. Alpha Diversity. Faith-pd diversity was used to compare community rarefaction curves (an area under the curve analysis) for both regions. Scrape sampling is shown to be generally lower in terms of diversity with tape sampling conversely the highest (p<0.0001).

Supplementary Figure 2. Nonmetric Multidimensional scaling (NMDS) of Bray-Curtis dissimilarity distance of skin microbiomes. The effect of sampling method (coloured dots) is shown for V1-3 (left panel) and V4 (right panel). Ellipses represent 95% CI. Tape sampling showed community composition to be statistically significantly different between tape and scrape for both V1-3 and V4, p=0.026 and p=0.002 respectively.

## Plain Language Summary

### Title: A detailed analysis of the best methods for analysis of skin bacteria

Healthy skin is covered with bacteria at a high density (approximately one million bacteria per square centimetre), which are known as the skin microbiome. The skin is used to having the bacteria on its surface and to some degree the bacteria in healthy skin are important in keeping the skin healthy. It is increasingly recognised that diseases affecting the skin often show an imbalance in the types and quantities of bacteria on the skin.

Historically, measurement of skin bacterial populations has relied on growth of the bacteria in a laboratory to quantities that become visible to the eye but is dependent upon the laboratory growing the bacteria efficiently. This approach has been superseded by a technique which does not rely on the bacteria growing and instead looks for the genetic signature of bacteria, and is therefore much better able to fully characterise the repertoire of bacteria from a sample. Because this new technology was largely developed in research on the gut microbiome, methodologies applied to skin, may not be appropriate.

Therefore, we set out to examine the skin microbiome using four sampling method (swabbing, scraping, swabbing after scraping and tape adhesion), as well as two different bacterial gene analysis approaches. Whilst many skin researchers have opted to follow the same protocol as gut analysis, we show that this is inferior for skin researchers. Additionally, we show that the optimal sampling method is tape adhesion. As a skin research community, we should harmonise these basic approaches to the study of skin microbiome, so that technical differences in methodology to not make it hard to compare different studies.

## Notes

**Declaration of Interests** None of the interests described below are considered to be conflicts with regard the work presented here. This work was carried out in partial fulfilment of RYA’s PhD thesis supported by the Royal Embassy of Saudi Arabia. DWC acknowledges support from Pfizer and GSK. MF was a consultant and member of the Scientific Advisory Board of AOBiome. MRAJ has been an adviser/consultant/speaker for AbbVie, Pfizer, Sanofi Genzyme, Lilly, Ducentis, Hosei Septares, Unilever.

### Competing Interest Statement

The authors have declared no competing interest.

### Summary of Updates

The 515F-806R primer pair described in the method only amplify V4 region NOT V3-4. Regards,

