## Supplementary Figure 1. for "Methods for the analysis of skin microbiomes: a comparison of sampling processes and 16S rRNA hypervariable regions"

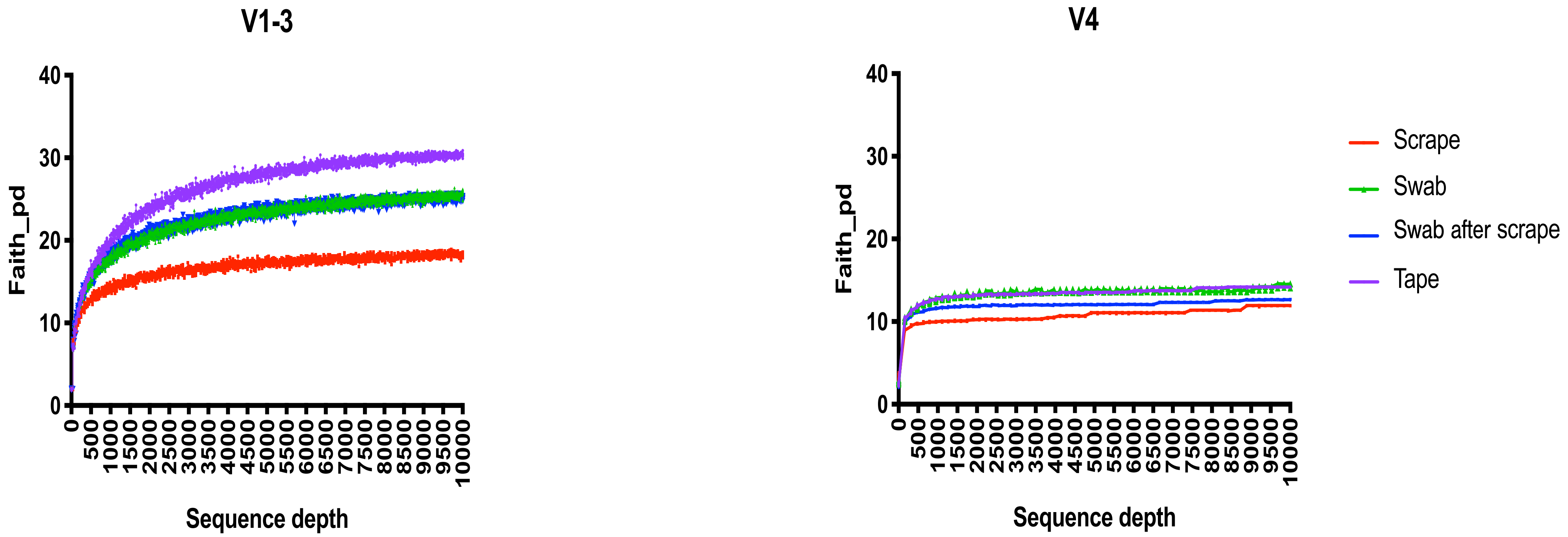


Figure 1. Alpha Diversity. Faith-pd diversity was used to compare community compositions for both regions showed that although all three approaches were better than scrape sampling, tape sampling was superior (p<0.0001).
