## Supplementary Figure 2. for "Methods for the analysis of skin microbiomes: a comparison of sampling processes and 16S rRNA hypervariable regions"

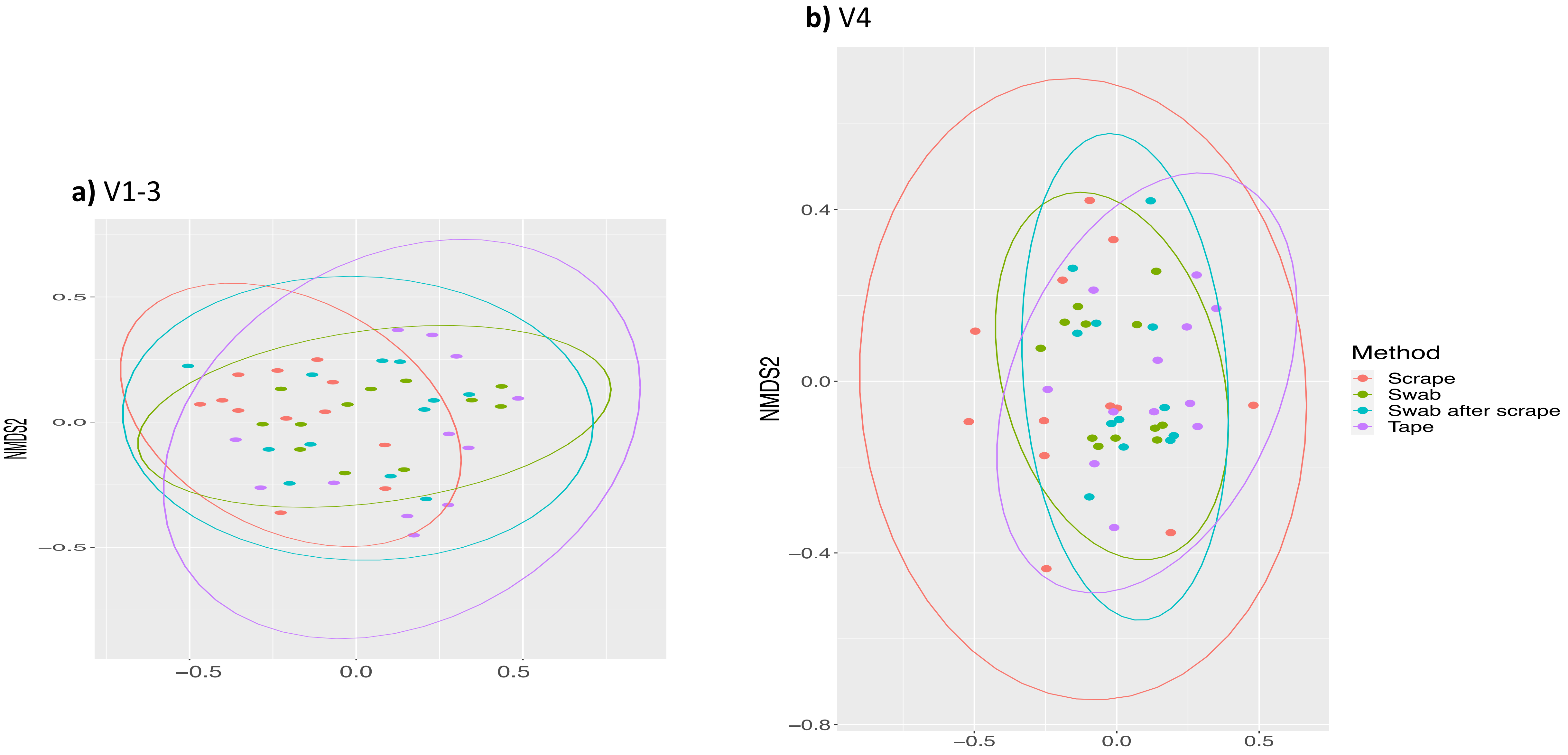


**Supplementary Figure 2.** Nonmetric Multidimensional scaling (NMDS) of Bray-Curtis dissimilarity distance of skin microbiomes. The effect of sampling method (coloured dots) is shown for V1-3 (left panel) and V4 (right panel). Ellipses represent 95% CI. Tape sampling showed community composition to be statistically significantly different between tape and scrape for both V1-3 and V4, p=0.026 and p=0.002 respectively.
